# Reassessment of lesion-associated gene and variant pathogenicity in focal human epilepsies

**DOI:** 10.1101/130203

**Authors:** Lisa M. Neupert, Michael Nothnagel, Patrick May, Aarno Palotie, Mark Daly, Peter Nürnberg, Ingmar Blümcke, Dennis Lal

**Author notes:** Corresponding Author: Dennis Lal, PhD, Cologne Center for Genomics, University of Cologne, Stanley Center for Psychiatric Research, Broad Institute, Cambridge, MA, Psychiatric & Neurodevelopmental Genetics Unit, Harvard Medical School / Massachusetts General Hospital Boston, MA, USA.

## Abstract

**Purpose:** Increasing availability of surgically resected brain tissue from Focal Cortical Dysplasia and low-grade epilepsy-associated tumor patients fostered large-scale genetic examination. However, assessment of germline and somatic variant pathogenicity remains difficult.

**Methods:** Here, we critically reevaluated the pathogenicity for all neuropathology-associated variants reported to date in the PubMed and ClinVar databases, including 12 disease-related genes and 88 neuropathology-associated missense variants. We (1) assessed evolutionary gene constraint using the *pLI* and missense *z* scores, (2) applied guidelines by the American College of Medical Genetics and Genomics (ACMG), and (3) predicted pathogenicity by using PolyPhen-2, CADD, and GERP.

**Results:** Constraint analysis classified only seven out of 12 genes to be likely disease-associated, while 35 (40%) of those 88 variants were classified as being variants of unknown significance (VUS) and 53 (60%) as being likely pathogenic (LPII). Pathogenicity prediction yielded discrimination between neuropathology-associated variants (LPII and VUS) and rare variant scores obtained from individuals present in the Genome Aggregation Database (gnomAD).

**Conclusion:** We conclude that several VUS are likely disease-associated and will be reclassified by future molecular evidence. In summary, interpretation of lesion-associated gene variants remains complex while the application of current ACMG guidelines including bioinformatic pathogenicity prediction will help improving interpretation and prediction.

## INTRODUCTION

Somatic gene variants have increasingly been detected and reported in brain tissue obtained from patients with epilepsy-associated focal lesions and considered causal for the lesion and the patient’s epilepsy. Most common structural brain lesions comprise Focal Cortical Dysplasia (FCD) [1, 2] and low-grade epilepsy associated tumors (LEAT) [3], both of which represent umbrella terms for a variety of diagnostically related but histologically independent etiologies. FCD are a heterogeneous group of cortical malformations accounting for the most common structural brain lesions within the broad spectrum of malformations of cortical development [4, 5]. FCD is diagnosed in 29-39% of patients who undergo epilepsy surgery [1], mostly affect the frontal lobe, and can histopathologically present with a large spectrum of abnormalities, including cortical architecture, bizarre neuronal cell morphology, blurred gray-white matter boundaries, and heterotopic neurons or increased oligodendroglial cell densities in white matter [5]. The most frequent tumors in patients with drug-resistant focal epilepsy starting in the first two decades of life are ganglioglioma (GG) and dysembryoplastic neuroepithelial tumors (DNT), representing 65% of 1,551 tumors collected at the European Epilepsy Brain Bank [6]. Intriguingly, these tumors histopathologically present with a variable mixture of glial and neuronal cell morphologies, and were mostly affecting the temporal lobe [6].

Unlike recent large-scale studies on rare and common germline variant-associated epilepsies, genetic studies on focal brain lesions comprise so far only small cohorts with insufficient patient numbers to perform reasonably powered hypothesis-free exome-wide gene discovery screens. Patient ascertainment is challenging since the disease-associated variants are expected to be present only in the brain or even in a fraction of the lesional brain tissue [1]. In addition to the limited access to the target tissue, the prevalence of somatic mosaicism in the human brain of healthy individuals is not well understood, and thus current small cohort studies without large control sets might lead to biased or even false conclusions. Even for genuine disease-associated genes, not all observed variants would contribute to disease etiology. Due to time and cost constrains, functional testing can usually be conducted only for a minor part of variants observed in patients. The latter represents a more general problem that holds true for all epilepsies since the accurate interpretation of variation in disease genes has largely lagged behind the massive up-scaling of data generation enabled by the increased accessibility of sequencing.

Here, we critically reevaluate the pathogenicity of genes and variants that have previously been reported to be associated with histopathologically confirmed brain lesions. We hypothesize that somatic variants in disease-associated genes, being present only in a small fraction of cells in the brain but leading to severe neuropathologies, should have substantially different molecular functional, evolutionary and prediction tool signatures when compared to rare benign germline variants that are located within the same genes and that are found in individuals from the general population in the Genome Aggregation database (gnomAD; http://gnomad.broadinstitute.org). The evaluation procedure represents state of the art evaluation of all reported neuropathology-associated as well as other disease-associated risk genes and variants and can be reapplied to other variant sets.

## MATERIALS AND METHODS

### Gene and variant identification

The evaluation of pathogenicity for loss-of-function variants (e.g. full gene deletions or nonsense variants) is relatively straightforward, however, for missense variants the criteria to establish pathogenicity rely on supportive genetic data and functional evidence [7]. Therefore, we focused on the interpretation of heterozygous dominant acting missense neuropathology-associated variants. Firstly, we performed a PubMed-based (https://www.ncbi.nlm.nih.gov/pubmed; accessed February 2017) literature review to identify studies that report genes and dominant acting missense neuropathology-associated variants. We used either single search terms or two-or three-word combinations of the following keywords: ‘focal cortical dysplasia’, ‘ganglioglioma’, ‘dysembryoplastic neuroepithelial tumor’, ‘neuropathology’, ‘genetics’, ‘somatic’ and ‘mutations’. Secondly, we collected all missense variants in genes associated with neuropathologies in the literature from the ClinVar database (http://www.ncbi.nlm.nih.gov/clinvar; release: 01/11/2016) using filters described in the **Supplementary Methods**. We removed copy number, synonymous, frameshift, splice site and nonsense variants from the dataset.

### Variant classification

All identified neuropathology-associated variants were classified in accordance to 31 criteria defined by guidelines of the The American College of Medical Genetics and Genomics (ACMG) [7]. We used the online tool provided by Kleinberger et al. [8] to aid the variant interpretation process. This tool displays all evidence categories from the ACMG guidelines and offers the possibility to select the appropriate criteria with simple checkboxes. We used the algorithm incorporated in the tool to assign either pathogenicity or a benign, non-deleterious impact based on the selected evidence categories, resulting in two positive variant groups in our study: 1) Variants of unknown significance (“VUS”) and 2) likely pathogenic variants (“LPII”).

In addition to the database-reported neuropathology-associated variants, we included two other groups in the pathogenicity prediction analysis as positive and negative comparison groups to guide the interpretation of our results with confidence 3) Functional variants from LEAT-associated genes that were also associated with non-brain tumors and had been validated in molecular studies [9]; ACMG guidelines categorized these variants as likely pathogenic variants (“LPIIC”). 4) Singleton germline variants in neuropathology-associated genes identified in >130k individuals from the general population (“GP”) from the Genome Aggregation Database (gnomAD; http://gnomad.broadinstitute.org; accessed November 2016).

### Assessment of evolutionary gene constraints

We evaluated literature-reported neuropathology-associated genes for evolutionary constraints by use of the *pLI* and missense *z* conservation scores [10] and compared the results with the proposed pathomechanism in the recent literature. These scores use a depletion of variants in a gene when compared to the expectation under neutral evolution. The expectation has been estimated from a population reference cohort of >60,000 individuals [10] as an indication of purifying selection, rendering variants affecting these genes more likely to be implicated in disease etiology. Following the authors’ recommendations, we considered genes with *pLI* scores >0.9 as being intolerant for loss-of-function (LoF) mutations and those with *z*-scores >3 intolerant for missense mutations.

### *In silico* evaluation of variant pathogenicity

We functionally annotated all missense neuropathology-associated (“VUS” and “LPII”), non-brain tumor-associated (“LPIIC”) and gnomAD singleton (“GP”) variants using three commonly used *in silico* prediction tools, namely CADD [11], GERP ++ [12] and PolyPhen-2 [13] with wANNOVAR [14]. Whereas CADD and PolyPhen-2 represent meta-tools, which incorporate multiple different biological and evolutionary scores, GERP ++ only scores conservation across species.

In order to identify the most accurate predictor to be used in subsequent analyses for distinguishing “LPII” and “VUS” variants from neutral ones (“GP”), we assessed predictor performance using the area under (AUC) receiver-operating-characteristic curves (ROC curves), thereby simultaneously assessing specificity and sensitivity, as implemented in the R package pROC (https://cran.R-project.org/package=pROC).

### Statistical analyses

Association *p*-values were obtained from a Wilcoxon ranked sum test, as implemented in the wilcox.test function of the R statistical software, version 3.3.1. [15]. A two-sided p-value < 0.05 was considered statistical significant. P-values were Bonferroni corrected in multiple testing. Furthermore, in order to correct for the effect of largely disparate group sizes, we performed 1000 times subsampling.

## RESULTS

### Assessment of evolutionary gene constraints

In our literature and ClinVar review, we identified a total of 12 genes associated with heterozygous dominant acting neuropathology-associated variants, namely *AKT3, BRAF, DEPDC5, FGFR1, IDH1, MTOR, NPRL2, NPRL3, PIK3CA, PTEN, TSC1, TSC2,* in 13 studies (Table S1). Six of 13 studies (46.2%) used whole exome sequencing (WES) as variant discovery technique, five of these combined with targeted sequencing or Sanger sequencing to identify variants in the rest of the cohort (Table S2). In total nine studies performed targeted sequencing of 1-14 genes (Median = 1, SD = 5.1). Only six out of 13 WES and targeted sequencing studies (46.2%) list detailed variant calling parameters and only six (46.2%) list all variants passing their filter criteria.

Based on the literature, we categorized *TSC1* and *PTEN* as intolerant for LoF, *MTOR*, *PIK3CA*, *BRAF*, *IDH1*, and *AKT3* intolerant for missense and *NPRL2*, *NPRL3*, *TSC2*, *FGFR1,* and *DEPDC5* as intolerant for both (Table 1). Conservation scores (*pLI* and missense *z)* indicated that only seven (*AKT3, BRAF, DEPDC5, MTOR, PIK3CA, PTEN, TSC1*) out of 12 genes showed support for association with a severe disease by harboring less variants than expected under neutral evolution.

**Table 1.** Evolutionary constraint for genes under investigation. Depletion score analysis of 12 neuropathology-associated genes according to the reported pathomechanism in the literature. ✔ = intolerant for reported mutations (*pLI* ≥ 0.9; *z* ≥ 3.0); X = tolerant for reported mutation (*pLI*<0.9, *z*<3.); “Yes” = reported pathomechanism with functional support; “No” = reported pathomechanism without functional support; *genes were excluded from the following evaluation of the variant pathogenicity.

| Gene | LoF intolerance in controls | Functional support |
| --- | --- | --- |
| PTEN | ✓ (0.98) | Yes |
| TSC1 | ✓ (1) | Yes |
| **Missense intolerance in controls** |  |  |
| AKT3 | ✓ (3.95) | Yes |
| BRAF | ✓ (3.99) | Yes |
| MTOR | ✓ (7.89) | Yes |
| PIK3CA | ✓ (5.42) | Yes |
| IDH1* | X (0.1) | No |
| **LoF/ Missense intolerance in controls** |  |  |
| DEPDC5* | ✓/✓ (1/3.29) | Yes |
| FGFR1 | ✓/ X (0.99/2.8) | Yes |
| TSC2 | ✓/ X (1/0.89) | Yes |
| NPRL2 | X/X (0.35/1.86) | No |
| NPRL3 | X/X (0.47/0.37) | No |

For ten out of those 12 neuropathology-implicated genes, disease-associated missense variants had been reported in patients. We next classified these variants according to recently published ACMG guidelines. Our literature and database review identified in total 88 neuropathology-associated missense variants (Figure 1a; Figure S1; Figure S2). Out of these, 53 (located in the *AKT3*, *BRAF*, *FGFR1, MTOR, PIK3CA* and *PTEN* genes) were classified as ‘likely pathogenic II (LPII)’, because they held several of the following criteria: showed a damaging effect on the gene or gene product in functional studies, were located in a mutational hot spot and/or critical and well-established functional domain, were absent from controls, had received a missense z score ≥ 3.0, showed a deleterious effect in multiple lines of computational evidence and/or were reported as pathogenic in a reputable source (Table S3). The ‘variant of uncertain significance (VUS)’ group comprised 35 variants present in all 10 neuropathology-associated genes, representing 1/3 of identified neuropathology-associated missense variants.

**Figure 1.**
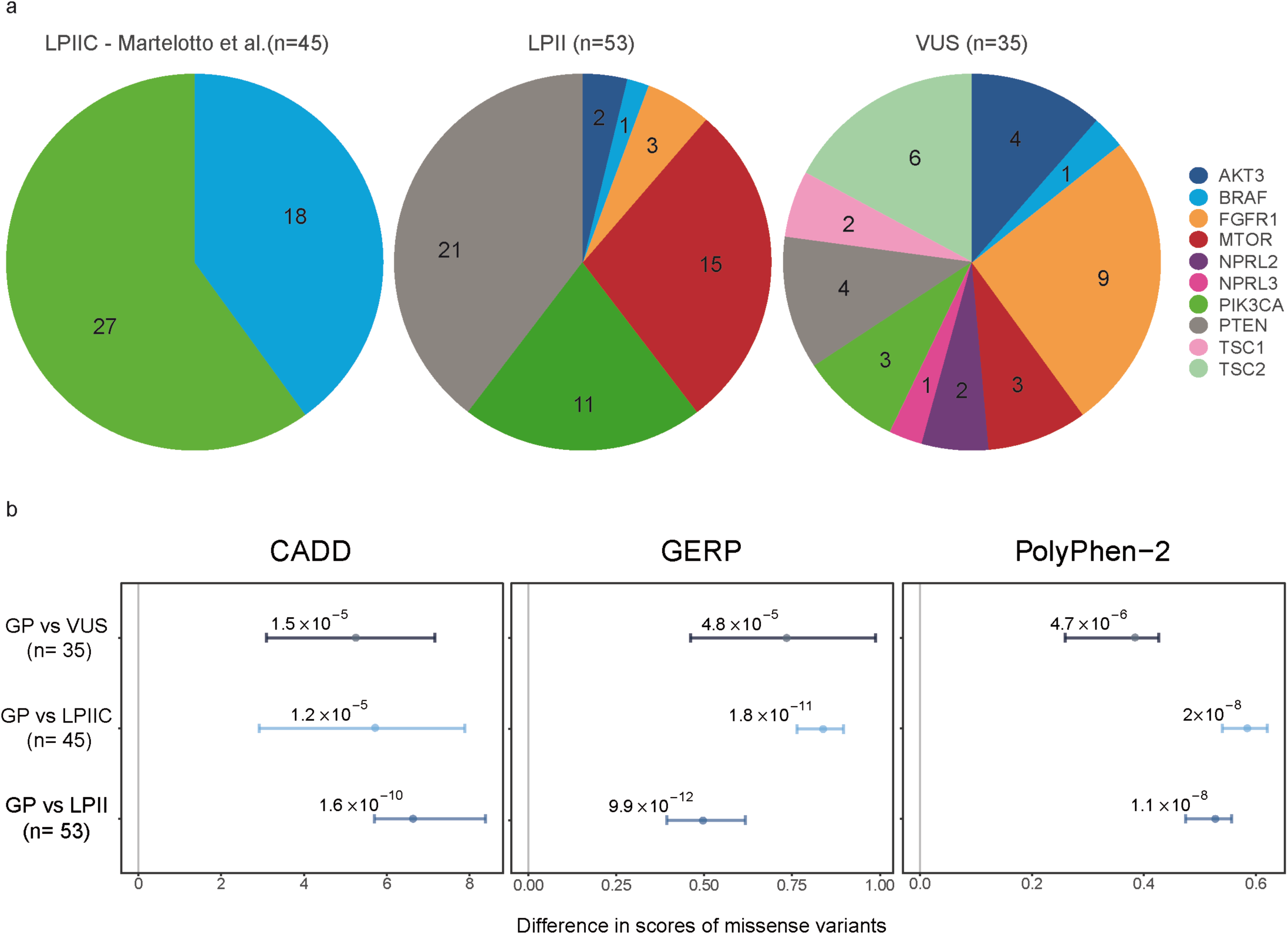
ACMG criteria classify neuropathology-associated and non-brain tumor-associated variants as being VUS or as being LPII variants. (**a**) Amounts of non-brain tumor-associated likely pathogenic variants (LPIIC), neuropathology-associated likely pathogenic variants (LPII), and neuropathology-associated variants of uncertain significance (VUS). The different colors indicate different neuropathology-associated genes. (**b**) Distribution of CADD, GERP and PolyPhen-2 missense variant predictions scores of groups VUS, LPIIC and LPII in comparison with rare variants from gnomAD (GP). P-values ≤ 0.017, Bonferroni corrected, were considered to be significant. CIs are indicated.

### *In silico* evaluation of variant pathogenicity

VUS categorization does not rule out disease association. It is quite conceivable that at least some of the 35 VUS truly might contribute to disease but currently not enough evidence has been generated for being classified as LPII. We explored this assumption further and investigated whether VUS variants behave *in silico* similar to LPII and known disease variants. We compared the missense variant prediction scores of functionally validated cancer variants, LPIIC variants (N=45, as positive control), the classified neuropathology-associated LPII (N=53) and of VUS (N=35) variants in comparison with rare variants from the general population (GP, as negative control) (Figure 1b). VUS, LPII and more pronounced LPIIC variants were significantly more pathogenic predicted than variants from the general population for in all prediction tool analyses (p ≤ 0.017, Bonferroni corrected; Figure 1b).

## DISCUSSION

We performed a comprehensive literature review to identify genes and missense variants associated with epilepsy-associated focal brain lesions. Evaluation of disease association using recently developed methods and guidelines confirmed pathogenicity for only 7 out of reported 12 genes and 53 (60%) out of reported 88 variants. Large variability observed in published assessment strategies challenges overall evidence and call for consensus use of international standards.

The majority of variants reported in neuropathology patients in previous studies were identified by targeted sequencing. In addition, detailed filtering parameters and full lists of identified variants are reported in the minority of studies. Accordingly, previous neuropathology studies reporting the identification of new disease-associated genes and variants were *a priori* hypotheses based and not unbiased.

Five of the 12 tested genes have yet insufficient support for classification as disease gene when investigating the evolutionary gene constraints. For example, dominant acting LoF variants in *NPRL2* and *NPRL3* have been reported in neuropathology patients [16] and, correspondingly, haploinsufficiency as pathomechanism has been proposed. The hallmark of haploinsuffient genes is the absence of LoF variants in healthy individuals [10]. However, *NPRL2* and *NPRL3*-affecting LoF variants have been reported in unaffected individuals and for both genes no statistically significant depletion of variants has been observed in large-scale database of healthy individuals from the general population. In addition, at least two papers reported germline missense, splicing and LoF variants in *NPRL3* in patients with FCDs and even showed nonsense-mediated decay for one variant present in a patient. Proof for the association of these genes with the disease will require, however, further statistical enrichment and *in vivo* modeling.

The prediction of pathogenicity missense variants is challenging and relies on next generation sequencing data and functional evidence [7]. For the 1/3 of evaluated variants, however, well-established *in vitro* or *in vivo* functional tests were not performed in patient tissue. Following the ACMG guidelines, such variants have to be classified as VUS if not compensated by other strong evidences. We did not find reliable evidence for 35 variants to be classified as likely pathogenic by working through all 31 ACMG classification criteria. However, our bioinfomatic prediction analysis showed that variant scores for VUS significantly deviate from those of rare variants from the general population, indicating that probably at least a fraction of these variants is truly associated with disease. In total, 6 of the VUS variants and 4 of the LPII variants were present in gnomAD and 50% of these occurred more than once in the general population whereas all 4 LPII variants were only singletons in gnomAD. Expecting that the identified patient variants act dominantly with large effect, their presence in general-population germline-variant databases like gnomAD suggested that these variants are unlikely to be associated with severe brain lesions occurring usually in early childhood [10].

Tremendous advances in sequencing technologies foster variant discovery at accelerating pace whereas translation into clinical variant classification remains in its infancy. This is particularly true in the neurosciences and for many brain diseases since variability of somatic mosaicism critically results from the diversity and admixture of neuroepithelial cell types and their developmental stage in a given brain tissue across humans. Based on our presented data, diagnostically relevant interpretation for novel missense variants is feasible only for a part of variants at this point. Results from large-scale projects, e.g. the human cell atlas and high throughput mutagenesis screening, will improve variant interpretation in the near future with consensus standards for assessment and interpretation widely implemented and used.

## SUPPLEMENTARY MATERIAL

Supplementary information is available at the *Genetics in Medicine* website.

## REFERENCES

1. Lim, J.S., et al., Brain somatic mutations in MTOR cause focal cortical dysplasia type II leading to intractable epilepsy. Nat Med, 2015. 21(4): p. 395–400.

2. Scheffer, I.E., et al., Mutations in mammalian target of rapamycin regulator DEPDC5 cause focal epilepsy with brain malformations. Ann Neurol, 2014. 75(5): p. 782–7.

3. Martinoni, M., et al., BRAF V600E mutation in neocortical posterior temporal epileptogenic gangliogliomas. J Clin Neurosci, 2015. 22(8): p. 1250–3.

4. Barkovich, A.J., et al., A developmental and genetic classification for malformations of cortical development: update 2012. Brain, 2012. 135(Pt 5): p. 1348–69.

5. Blumcke, I. and R. Spreafico, An international consensus classification for focal cortical dysplasias. Lancet Neurol, 2011. 10(1): p. 26–7.

6. Blumcke, I., et al., A neuropathology-based approach to epilepsy surgery in brain tumors and proposal for a new terminology use for long-term epilepsy-associated brain tumors. Acta Neuropathol, 2014. 128(1): p. 39–54.

7. Richards, S., et al., Standards and guidelines for the interpretation of sequence variants: a joint consensus recommendation of the American College of Medical Genetics and Genomics and the Association for Molecular Pathology. Genet Med, 2015. 17(5): p. 405–24.

8. Kleinberger, J., et al., An openly available online tool for implementing the ACMG/AMP standards and guidelines for the interpretation of sequence variants. Genet Med, 2016. 18(11): p. 1165.

9. Martelotto, L.G., et al., Benchmarking mutation effect prediction algorithms using functionally validated cancer-related missense mutations. Genome Biol, 2014. 15(10): p. 484.

10. Lek, M., et al., Analysis of protein-coding genetic variation in 60,706 humans. Nature, 2016. 536(7616): p. 285–91.

11. Kircher, M., et al., A general framework for estimating the relative pathogenicity of human genetic variants. Nat Genet, 2014. 46(3): p. 310–5.

12. Davydov, E.V., et al., Identifying a high fraction of the human genome to be under selective constraint using GERP++. PLoS Comput Biol, 2010. 6(12): p. e1001025.

13. Adzhubei, I.A., et al., A method and server for predicting damaging missense mutations. Nat Methods, 2010. 7(4): p. 248–9.

14. Yang, H. and K. Wang, Genomic variant annotation and prioritization with ANNOVAR and wANNOVAR. Nat Protoc, 2015. 10(10): p. 1556–66.

15. R Core Team. R: A Language and Environment for Statistical Computing. 2016 [cited 2015; Available from: http://www.R-project.org/.

16. Weckhuysen, S., et al., Involvement of GATOR complex genes in familial focal epilepsies and focal cortical dysplasia. Epilepsia, 2016. 57(6): p. 994–1003.

